# Cryptic adaptor protein interactions regulate DNA replication initiation

**DOI:** 10.1101/313882

**Authors:** Lindsay A. Matthews, Lyle A. Simmons

## Abstract

DNA replication is a fundamental biological process that is tightly regulated in all living cells. In bacteria, the master regulator DnaA controls when and where replication begins by building a step-wise complex that loads the replicative helicase onto chromosomal DNA. In many bacteria, DnaA requires the adaptor proteins DnaD and DnaB to aid DnaA during helicase loading. How DnaA, its adaptors, and the helicase form a complex at the origin is largely unknown. In this study, we addressed this long-standing question by disassembling the initiation proteins into their individual domains and testing all possible pair-wise combinations in a bacterial two-hybrid assay. Here we report a full description of the cryptic interaction sites used by the helicase loading machinery from *Bacillus subtilis*. In addition, we investigated how complex formation of the helicase loading machinery is regulated by the checkpoint protein SirA, which is a potent replication inhibitor in sporulating cells. We found that SirA and the DnaD adaptor bind overlapping sites on DnaA, and therefore SirA acts as a competitive inhibitor to block initiation. The interaction between DnaA and DnaD was also mapped to the same DnaA surface in the human pathogen *Staphylococcus aureus*, demonstrating the broad conservation of this interface. Therefore, our approach has unveiled key protein interactions essential for initiation and is widely applicable for mapping interactions in other signaling pathways that are governed by cryptic binding surfaces.

**Author Summary:** In order to proliferate, bacteria must first build a step-wise protein complex on their chromosomes that determines when and where DNA replication begins. This protein complex is assembled through dynamic interactions that have been difficult to study and remain largely uncharacterized. Here we show that by deconstructing the proteins into their constituent domains, the interactions used to build the initiation complex can be readily detected and mapped to single amino acid resolution. Using this approach, we demonstrate that DNA replication is controlled through conformational changes that dictate the availability of interaction surfaces. In addition, negative regulators can also block DNA replication by influencing complex formation so that cells survive inhospitable conditions. Initiation proteins from the model organism *B. subtilis* and the human pathogen *S. aureus* were both used to underscore the general applicability of the results to different bacterial systems. Furthermore, our general strategy for mapping dynamic protein interactions is suitable for many different signaling pathways that are controlled through cryptic interaction surfaces.

## Introduction

All cells use a dedicated replicative helicase loading pathway to ensure DNA replication begins at the right time and place. Bacteria use the master regulator DnaA to recognize the origin of replication (*oriC*) and induce melting of the origin DNA (1). DnaA then recruits the replicative helicase with its associated loader, representing the first step in replication fork assembly. DnaA is widely conserved, but importantly its mechanism differs between Gram-positive and Gram-negative species. In *Escherichia coli*, DnaA recruits the replicative helicase through a direct interaction (2,3). In contrast, the Gram-positive model organism *B. subtilis* requires the essential adaptor proteins DnaD and DnaB to allow for helicase loading. Specifically, DnaA recruits DnaD to the origin first; DnaD then recruits DnaB; and, finally, DnaB recruits and helps load the helicase (4).

Interactions involving the DnaD and DnaB adaptors are tightly regulated and therefore add a layer of control over origin firing that is not present in Gram-negative systems. For example, *B. subtilis* is able to sporulate as a means of surviving inhospitable conditions and DNA replication initiation must be prevented during sporulation. This inhibition is due to a small protein called SirA that is only produced once the cell is committed to sporulate (5,6). SirA directly binds to a hot spot on the DnaA initiator and acts as a potent inhibitor of its activity (7,8). Though the mechanism of action is unknown, SirA prevents DnaA from accumulating at the origin and recruiting the DnaD and DnaB adaptors (7).

DnaD and DnaB share similar protein architectures and are conserved among Gram-positive bacteria with low GC-content, which includes many human pathogens such as *Staphylococcus aureus*, *Listeria monocytogenes*, *Streptococcus pneumoniae*, and *Bacillus anthracis* (9). In addition to DnaA, other types of initiators also use DnaD and DnaB to load the replicative helicase. For example, PriA uses DnaD and DnaB to restart stalled replication forks and some plasmid initiators use DnaD and DnaB during plasmid duplication (10–12). Multi-drug resistance plasmids and phage genomes can also express their own versions of DnaD- and DnaB-like adaptor proteins when replicating in low GC-content Gram-positive hosts (13,14). Therefore, determining how DnaD and DnaB function is critical for understanding DNA replication in multiple contexts. To this end, we sought to determine if and how interactions mediated by the DnaD and DnaB adaptors are regulated.

Interactions involving the DnaD and DnaB adaptors have been difficult to study using traditional screening methods. The DnaA and DnaD interaction gives mixed results in yeast two-hybrid assays depending on the experimental conditions (15). In contrast, DnaD and DnaB do not generate any signals with each other in yeast two-hybrid assays (13,15,16). There is some evidence that DnaD and DnaB can weakly interact *in vitro* (12,17), though currently the strongest support of a direct interaction is a gain-of-function mutation in DnaB that generates a robust interaction with DnaD and increases the frequency of initiation *in vivo* (13). This gain-of-function mutation (S371P) has been mapped to the C-terminus of DnaB and likely induces a conformational change, though it is unclear if this mimics a physiological process. Consequently, there is a pressing need to establish a method for detecting interactions with the wild type DnaD and DnaB adaptors.

To investigate how the DnaD and DnaB adaptors control replication initiation, we used a bacterial two-hybrid (B2H) assay to map their interaction surfaces. While the full-length initiation proteins did not interact, specific combinations of isolated domains from DnaA, DnaD, DnaB, and the replicative helicase produced robust signals. Using this approach, the interaction between DnaA and DnaD was mapped to the same hot spot used by the SirA inhibitor. Therefore, we propose that SirA acts as a competitive inhibitor for the DnaD adaptor. In addition, we found that DnaD used its N-terminal winged helix domain to mediate all of its interactions, while DnaB instead favored its C-terminal domain to bind to its partners. Together, these results indicate that protein interactions at the *B. subtilis* origin are controlled through conformational changes that expose cryptic interaction surfaces to build a step-wise initiation complex.

## Results

### Identification of interaction sites in initiation proteins

A B2H assay was used to detect interactions between the *B. subtilis* initiation proteins (**Fig 1A**). Positive interactions generate cyclic AMP within a *cya-E. coli* host cell, which can be detected as blue colonies in the presence of X-Gal or growth using maltose as the only carbon source (18). Both DnaD and DnaB homo-oligomerize (12) and these self-interactions could be readily detected in the assay (**Fig 1B**). In contrast, DnaD and DnaB did not interact with each other or with DnaA under the same conditions (**Fig 1C**). As a further control, a DnaB variant with the S371P gain-of-function mutation was also used (13), which generated a signal within the B2H assay as expected (**Fig 1C**).

**Figure 1.**
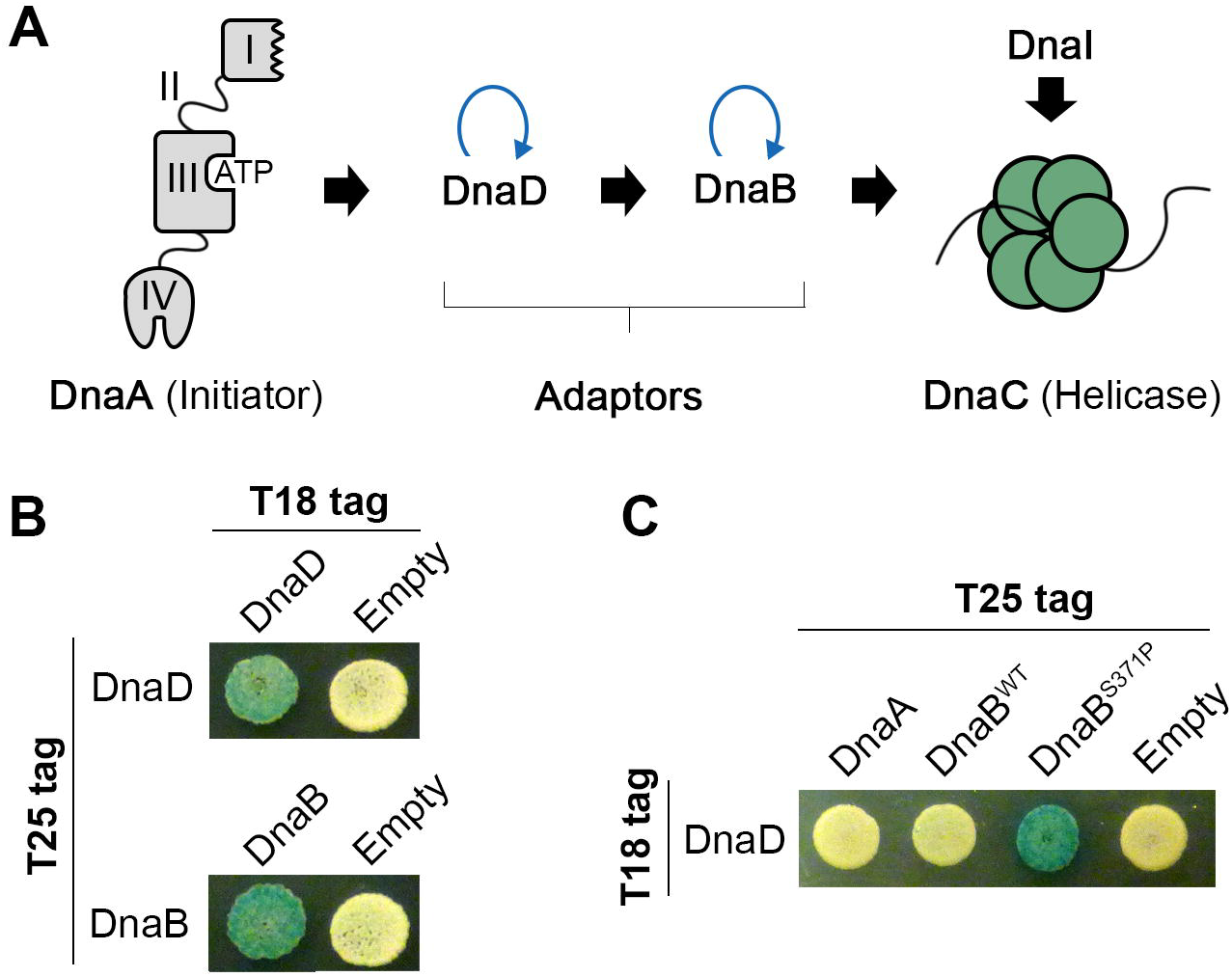
Full-length DnaA, DnaD and DnaB do not interact in a B2H assay. **(A)** A schematic of the helicase loading pathway in *B. subtilis*. Black arrows indicate potential protein-protein interactions while blue curved arrows indicate self-interactions. **(B)** B2H of T18- with T25-tagged DnaD (top) or DnaB (bottom) to demonstrate self-interactions mediated by the adaptor proteins. **(C)** B2H of T18-tagged DnaD co-expressed with either T25-tagged DnaA, wild type DnaB (DnaB^WT^), or the gain-of-function DnaB variant (DnaB^S371P^).

As we could not detect interactions between the full-length proteins, we chose an alternative strategy to map their interaction surfaces. DnaA, DnaD, and DnaB were separated into their constituent domains (**Fig 2A**) and tested against each other in all possible pairwise combinations using both X-Gal and minimal media containing maltose to determine positive interactions. This approach ensures that all potential binding sites hidden in the conformation favored by the full-length protein are exposed.

**Figure 2.**
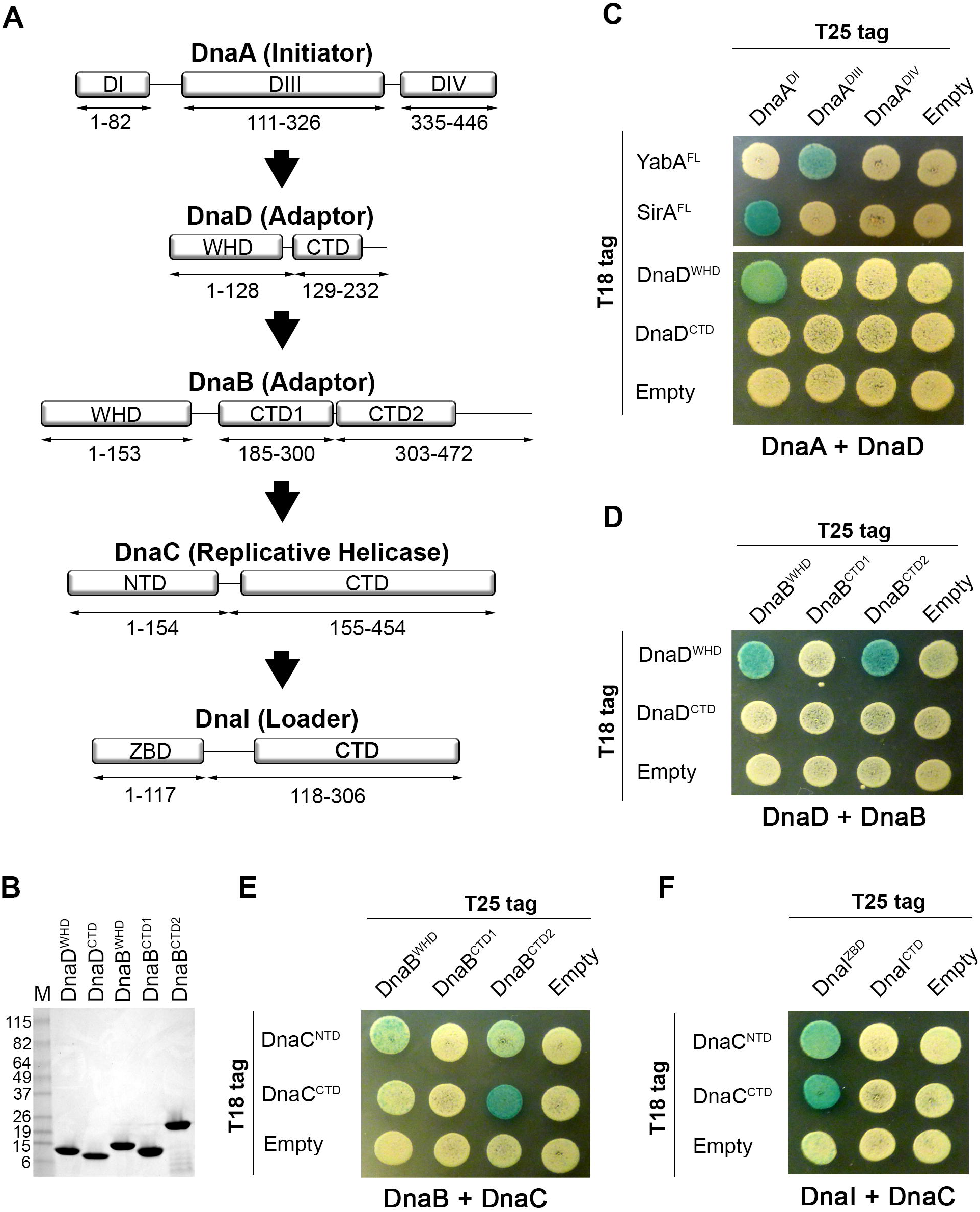
Mapping Interacting Domains Involved in DNA Replication Initiation. **(A)** Schematic of DnaA, DnaD, DnaB, DnaC, and DnaI divided into their individual domains. Amino acid boundaries of each protein fragment are indicated. WHD = Winged Helix Domain; CTD = C-terminal Domain; NTD = N-terminal Domain; ZBD = Zinc Binding Domain. **(B)** SDS-polyacrylamide gel showing purified domains from DnaD and DnaB. **(C)** B2H of T25-tagged DnaA domains co-expressed with T18-tagged full-length YabA (YabA^FL^), full-length SirA (SirA^FL^), or the domains from DnaD. **(D)** B2H of T25-tagged DnaB domains co-expressed with T18-tagged DnaD domains. **(E)** B2H of T25-tagged DnaB domains co-expressed with T18-tagged DnaC domains. **(F)** B2H of T25-tagged DnaI domains co-expressed with T18-tagged DnaC domains.

To test that the domain boundaries were correct, we purified each individual domain from the DnaD and DnaB adaptors and found they could fold as stable products (**Fig 2B**). For DnaA, we tested its domains with known binding partners in the B2H. DnaA domain I (DnaA^DI^) was found to interact with the SirA inhibitor (8) while DnaA domain III (DnaA^DIII^) interacted with YabA as expected (19) (**Fig 2C**). Therefore, the individual domains appear to be suitable for screening in the B2H assay as they are stable and exhibit other known functions.

For the DnaA-DnaD interaction, we found that the winged helix domain from DnaD (DnaD^WHD^) interacted with DnaA^DI^ (**Fig 2C**). DnaD^WHD^ also interacted with its downstream partner, DnaB, but in this case the signal was split between the DnaB winged helix domain (DnaB^WHD^) and its second C-terminal domain (DnaB^CTD2^) (**Fig 2D**). These results were also corroborated when using growth on minimal media containing maltose (**S1 Fig**). The winged helix domain of DnaD mediates a dimer-tetramer equilibrium; however, expressing DnaD^WHD^ on its own shifts the equilibrium to favor the tetrameric form (20). We entertained the possibility that the tetrameric form of DnaD^WHD^ specifically interacts with its protein partners. To test this directly, DnaD^WHD^ was truncated to prevent its tetramerization as determined by size exclusion chromatography, but this did not disturb any of the B2H signals (**S2 Fig**). Therefore, it is unlikely that the oligomeric state of DnaD dictates its interaction status. Instead, our results indicate that full-length DnaA, DnaD and DnaB may undergo conformational changes to expose cryptic binding sites and form stable complexes with each other during initiation.

The same B2H method was also used to map the interaction between the DnaB adaptor and the replicative helicase (called DnaC in *B. subtilis*; not to be confused with the DnaC loader from *E. coli*). The major signal came from DnaB^CTD2^ interacting with the C-terminal end of the helicase (DnaC^CTD^), though weak signals with DnaC^NTD^ were also apparent (**Fig 2E**). The zinc binding domain of the *B. subtilis* ATPase loader (called DnaI) is also known to interact with the C-terminal end of the DnaC helicase, but interactions between DnaI and DnaC^NTD^ have not been investigated (16, 21–23). Therefore, the DnaI domains (**Fig 2A**) were screened against the DnaC helicase domains. We found that the DnaI zinc binding domain (DnaI^ZBD^) can interact with both the N- and C-terminal ends of the helicase (**Fig 2F**). These results were also reproduced using growth on minimal media supplemented with maltose (**S3 Fig**). Therefore, we suggest that DnaI may have multiple interaction modes with the replicative helicase during loading.

### The DnaD wing interacts with the DnaA initiator

After identifying all of the interacting domains, we chose to focus on the interaction between DnaA and DnaD as this is likely the most regulated step of the helicase loading pathway. The ConSurf server was used to map conserved residues on the DnaD^WHD^ crystal structure [PDB 2V79 (20)] and reveal potential interaction sites (24–27). A conserved patch was found in a cleft formed by the β-strands of the wing in DnaD^WHD^ (**Fig 3A**). When the conserved hydrophobic cleft residues F51 and I83 were mutated to alanines the interaction with DnaA^DI^ was lost (**Fig 3B**). Furthermore, reducing the length of the loop (Δloop) in the DnaD^WHD^ wing (20) also disrupted binding to DnaA^DI^ (**Fig 3B**). This indicates that the DnaD wing region is necessary for forming a complex with DnaA. To ensure the F51A and I83A DnaD^WHD^ mutations did not cause structural defects, we purified both variants and verified that they could tetramerize using size exclusion chromatography (data not shown) and chemical cross-linking (**Fig 3C**). Tetramerization uses surfaces distal to the wing and is therefore an excellent test for wide-ranging structural defects (20). We also found that the DnaD^WHD^ wing variants could still interact with DnaB (**Fig 3B**), which further indicates they are structurally sound. Note that the DnaD^WHD^ Δloop variant was previously found to be stable and therefore was not tested (20).

**Figure 3.**
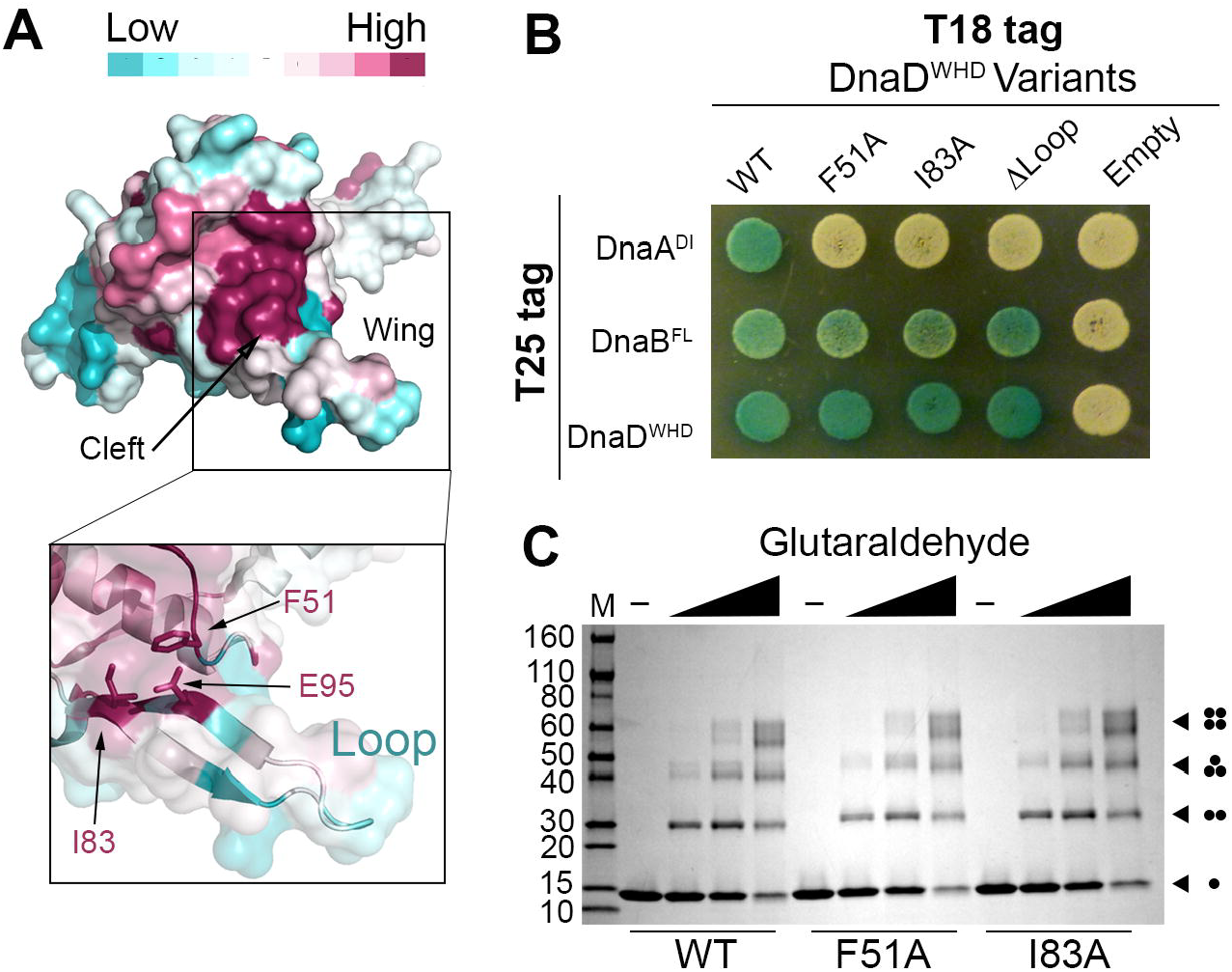
The DnaD^WHD^ Wing Forms a Binding Cleft for DnaA. **(A)** The crystal structure of DnaD^WHD^ (PDB 2V79) is shown as a surface representation and colored according to sequence conservation as indicated in the legend (top). The boxed inset shows a semi-transparent surface with F51, I83 and E95 represented as sticks. **(B)** B2H of T18-tagged DnaD^WHD^ variants co-expressed with either T25-tagged DnaA^DI^, DnaB^FL^ or DnaD^WHD^. **(C)** SDS-polyacrylamide gel stained with coomassie blue showing the glutaraldehyde cross-linking of wild type DnaD^WHD^ (WT) or the F51A and I83A variants to reveal self-interactions. The various oligomeric forms are symbolized by dots at the right-hand side of the gel, with each dot representing one DnaD^WHD^ protomer. The molecular weight marker is labeled on the left-hand side in kDa.

To test if the wing region is also critical in full-length DnaD, we used a temperature sensitive *B. subtilis* strain (*dnaD23*) where the native DnaD protein is not functional at 48°C (28). Ectopically expressing the F51A or I83A full-length DnaD variants from the *amyE* locus did not rescue this strain at the restrictive temperature (**Fig 4A**). Therefore, the DnaD wing is also required for function *in vivo*.

**Figure 4.**
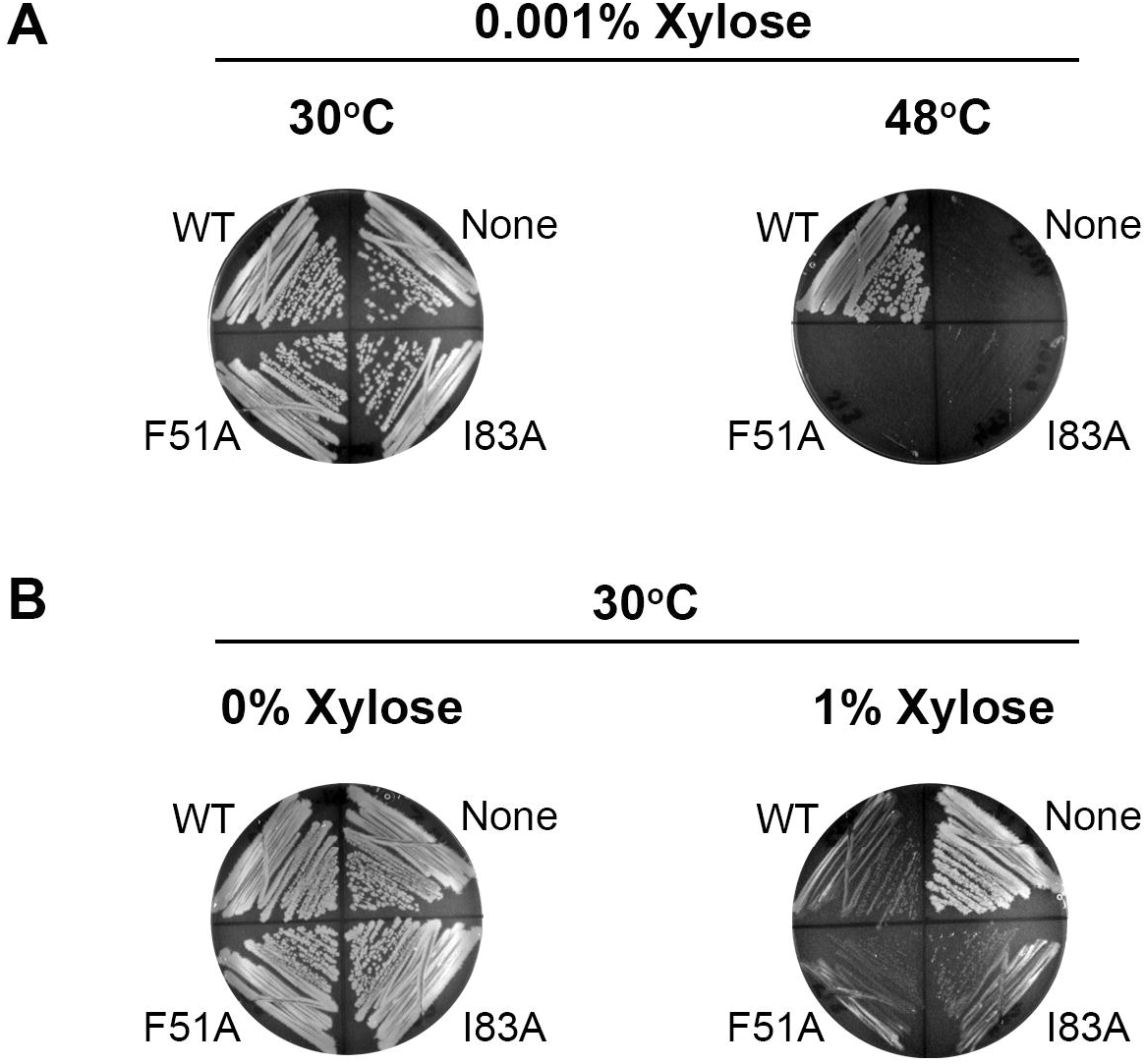
The DnaD^WHD^ Binding Cleft is Essential. **(A)***dnaD23* strains ectopically expressing WT DnaD or its winged helix variants (F51A or I83A) were incubated at the permissive temperature (30°C) or the restrictive temperature (48°C). The *dnaD23* parental strain was also included as a control (labeled “None” on the plates). **(B)** The same strains described in **(A)** were also incubated at the permissive temperature without (0%) or with (1%) xylose induction. Small colonies indicate a growth defect upon DnaD over-expression.

Overexpressing DnaD in *B. subtilis* causes a growth defect that manifests in small colony sizes. This phenotype depends on DNA binding by DnaD but is independent of its interaction with DnaA (9). Consequently, we predicted that if the F51A and I83A DnaD variants were only defective in binding DnaA, but not in other functions, they should also induce a small colony phenotype when overexpressed. The *dnaD23* strains used in Fig 4A were therefore incubated with high levels of xylose at the permissive temperature. Under these conditions, both F51A and I83A DnaD produced small colonies (**Fig 4B**), which suggests functions outside of DnaA binding are still preserved.

### DnaD and SirA bind the same interaction hot spot in DnaA

As SirA and DnaD both bind to DnaA^DI^, we asked whether they compete for the same binding site. SirA binds to an interaction hot spot on DnaA^DI^ that intriguingly cannot tolerate mutations *in vivo* despite the fact that the interaction with SirA is not essential (8). We hypothesized that DnaD may also bind to this DnaA^DI^ hot spot and that mutating this region is lethal because it knocks out the interaction with the DnaD adaptor.

To test our hypothesis, six different DnaA^DI^ variants were used: three with mutations on the hot spot that are lethal in *B. subtilis* (T26A, W27A, and F49A) (8) and three with mutations on the same surface that are still viable *in vivo* (N47A, F49Y, and A50V) (7). In the B2H assay, only the mutations that are lethal in *B. subtilis* knocked out the interaction with DnaD (**Fig 5A**). As a control, the DnaA variants were also tested against SirA where they all knocked out the interaction (**Fig 4D**), with the exception of T26A which is consistent with previous studies (8). Therefore, DnaD and SirA bind to overlapping surfaces on DnaA^DI^.

**Figure 5.**
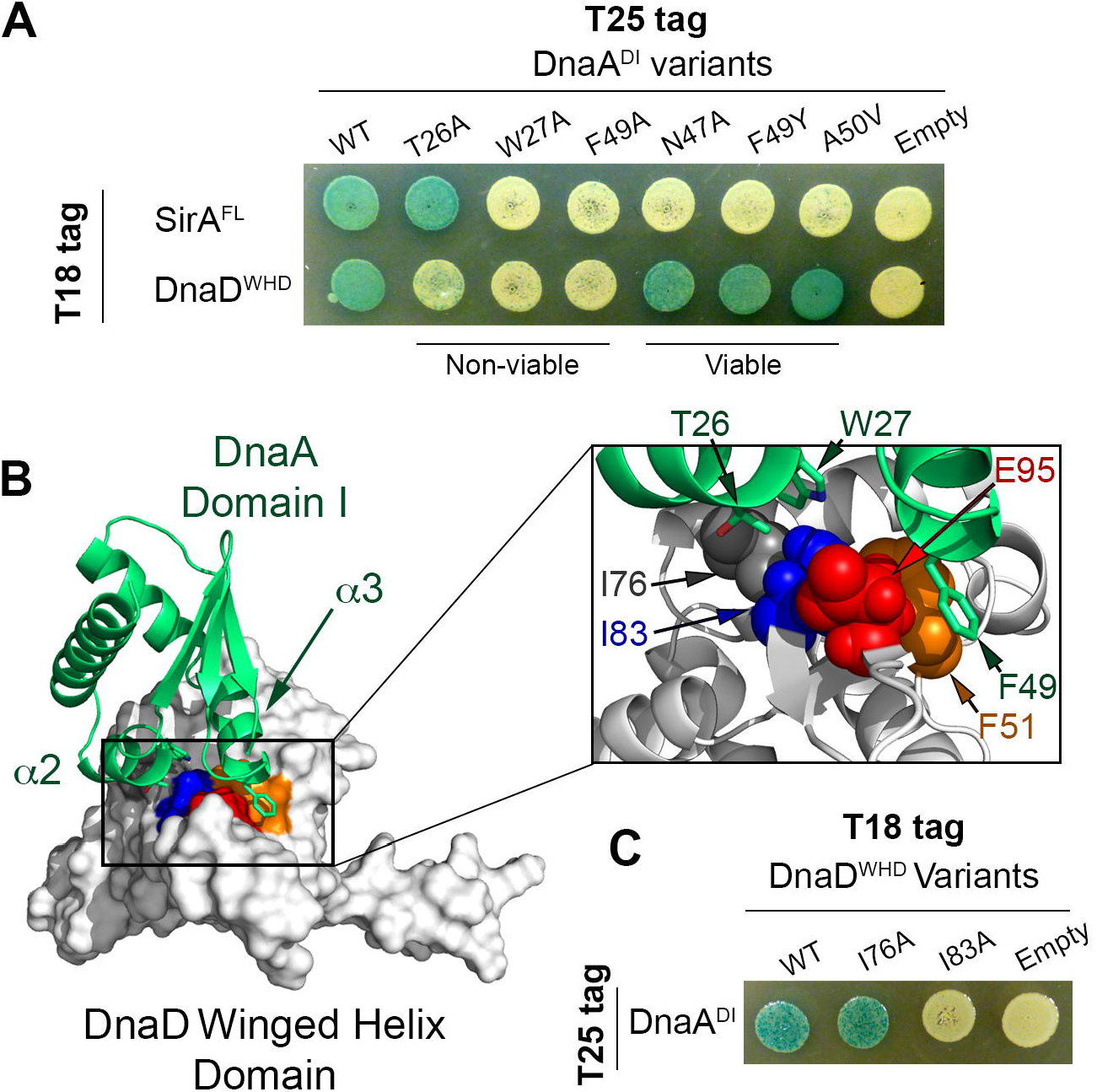
DnaD^WHD^ Binds to the DnaA^DI^ hot spot. **(A)** B2H of T25-tagged DnaA^DI^ variants co-expressed with T18-tagged SirA or DnaD^WHD^. “Viable” and “non-viable” refer to the effect these DnaA variants have when introduced into *B. subtilis*. **(B)** Model of the DnaD^WHD^ and DnaA^DI^ complex. DnaA^DI^ is represented as a green ribbon diagram with ***α***2 and ***α***3 labeled, while DnaD^WHD^ is shown as a white surface. The conserved residues that line the binding cleft in DnaD^WHD^ are colored with F51 in orange, I83 in blue, and E95 in red. The boxed inset shows a zoomed view of the interaction interface with the DnaD^WHD^ F51, I83 and E95 sidechains shown as spheres. DnaD^WHD^ I76 is also shown and colored in grey. The DnaA^DI^ interacting residues (T26, W27 and F49) are shown as green sticks with oxygen atoms colored red and nitrogen atoms colored blue. **(C)** B2H of T25-tagged DnaA^DI^ co-expressed with T18-tagged DnaD^WHD^ variants.

As crystal structures of both partners are available, we also modeled the complex between DnaD^WHD^ and DnaA^DI^ using the Rosetta docking algorithm (29–31). This model predicted a hydrophobic interaction surface on DnaD consisting of I83, F51 and the aliphatic portion of the E95 sidechain (**Fig 5B**). DnaA^DI^ docks on this hydrophobic surface using T26, W27 and F49 (**Fig 5B**). Therefore, the residues identified through our B2H assays are predicted to be in direct contact within the complex.

The DnaD crystal structure indicated that I83 may serve a structural role by packing against I76 and stabilizing the DnaD wing (**Fig 5B**). We entertained the possibility that mutating I83 increases the flexibility of the wing region by destroying this anchoring point; however, this is unlikely to be the reason the interaction with DnaA was lost as mutating the other isoleucine involved (I76A) did not prevent DnaA binding (**Fig 5C**). Therefore, we concluded that I83 is directly important for complex formation.

### *S. aureus* DnaA binds its cognate DnaD adaptor using the domain I hot spot

SirA and DnaD bind to overlapping surfaces on the DnaA domain I hot spot, which indicates these two factors compete with each other during early stages of sporulation. Interestingly, the domain I hot spot is widely conserved across DnaA homologs (32). This led us to ask if the DnaA hot spot also recruits DnaD to the origin in species that lack SirA, or whether these organisms instead evolved a different interaction site (**Fig 6A**).

**Figure 6.**
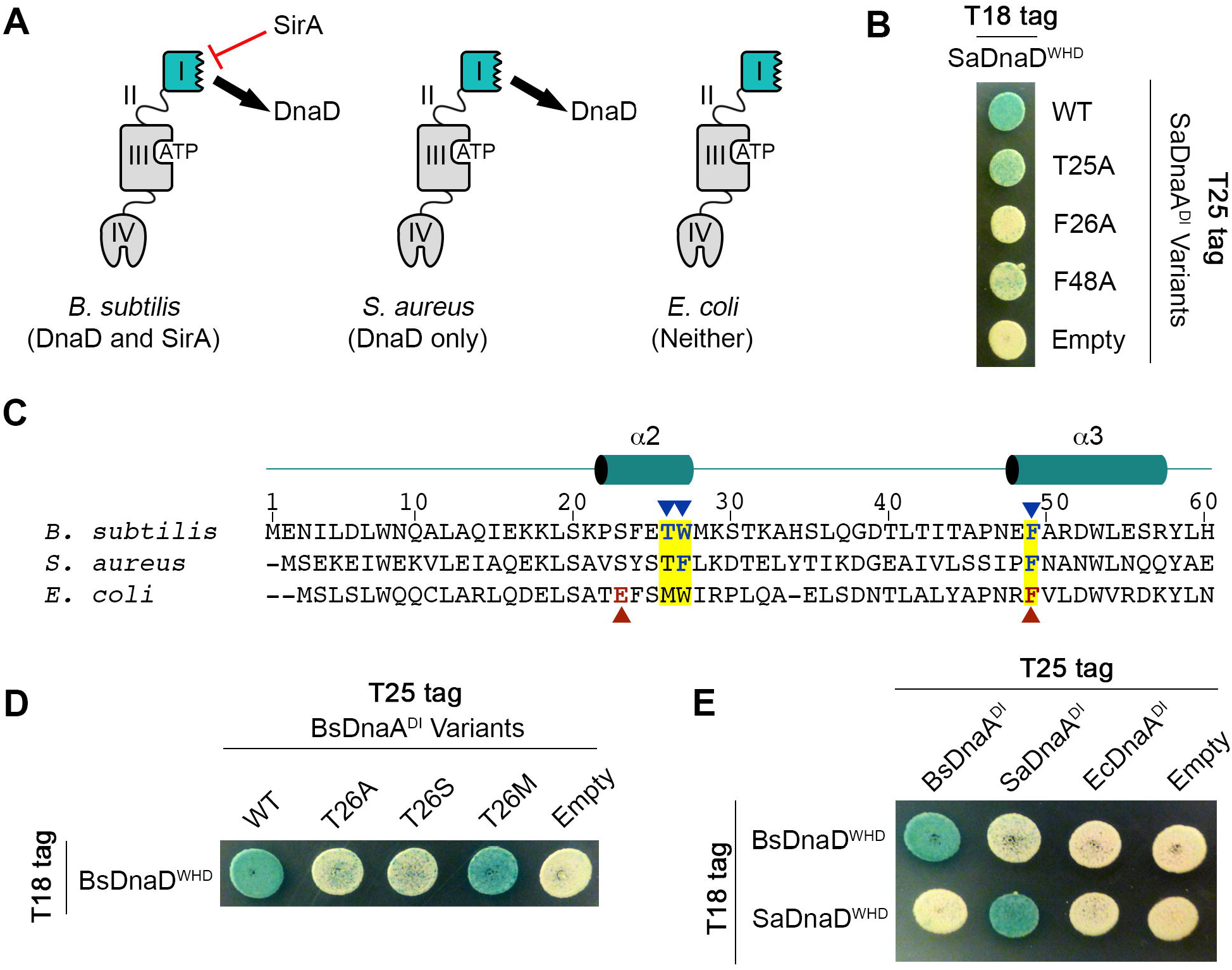
DnaD Binds to the DnaA domain I hot spot in *S. aureus*. **(A)** Schematic of DnaA demonstrating three different possibilities in bacteria: species that have both the DnaD adaptor and SirA regulator, species that have DnaD but lack SirA, and species that lack DnaD and SirA. **(B)** B2H of T18-tagged SaDnaD^WHD^ co-expressed with T25- tagged variants of SaDnaA^DI^. **(C)** Sequence alignment of the DnaA hot spot from *B. subtilis*, *S. aureus*, and *E. coli*. The three hot spot residues mutated in the B2H assays are highlighted in yellow, with residues critical for the DnaD interaction in *B. subtilis* indicated by blue arrow heads. Residues critical for the direct interaction between *E. coli* DnaA and the replicative helicase are indicated by red arrow heads. The numbering at the top of the alignment refers to the *B. subtilis* sequence. **(D)** B2H of T18-tagged BsDnaD^WHD^ variants co-expressed with T25-tagged BsDnaA^DI^. **(E)** B2H of T25-tagged DnaA^DI^ from *B. subtilis* (Bs), *S. aureus* (Sa) or *E. coli* (Ec) co-expressed with T18- tagged BsDnaD^WHD^ or SaDnaD^WHD^.

To test this possibility, we used the human pathogen *S. aureus* because it lacks a SirA homolog. DnaA and DnaD from *S. aureus* (herein referred to as SaDnaA and SaDnaD) interacted in a B2H using domain I from SaDnaA and the winged helix domain from SaDnaD (**Fig 6B**). The hot spot residues equivalent to T26, W27 and F49 in *B. subtilis* DnaA (T25, F26, and F48 in SaDnaA) were then mutated to alanines (**Fig 6C**). Mutating the two aromatic residues on the interaction hot spot (SaDnaA F26A and F48A) severely decreased the signal in the B2H assay, while mutating the threonine (SaDnaA T25A) had only a modest effect (**Fig 6B**). Therefore, the hot spot on DnaA domain I still interacts with its cognate adaptor in species that lack SirA, but the molecular contacts differ slightly from the complex in *B. subtilis*.

Out of the three residues on the DnaA hot spot investigated to this point, the contribution of the conserved threonine (T26 in BsDnaA; T25 in SaDnaA) was still unclear. Our model predicted that the interaction between DnaA and DnaD is primarily hydrophobic in *B. subtilis*, with the bulky methyl group from the threonine contributing directly to the interaction interface (**Fig 5B**). To test this further, T26 in BsDnaA was mutated to a serine which would effectively remove only the methyl group in question. This mutation is sufficient for knocking out the interaction with DnaD in a B2H (**Fig 6D**). To determine if other hydrophobic residues could substitute for T26 in the *B. subtilis* DnaA-DnaD complex, we also tested a T26M mutation and found the interaction with DnaD was maintained (**Fig 6D**). These results strongly support the model that the DnaA hot spot is providing a primarily hydrophobic surface that docks onto the DnaD wing. Therefore, it appears that the conserved threonine contributes to this hydrophobic hot spot in *B. subtilis* DnaA, but is largely dispensable in *S. aureus*.

The residues on the DnaA hot spot are highly conserved, yet they usually mediate species-specific contacts (32). To determine if the interaction with the DnaD adaptor is also species-specific, we tested if different DnaA homologs could bind the DnaD adaptor from either *B. subtilis* or *S. aureus* in a B2H. The results demonstrated that both BsDnaA and SaDnaA could only recognize their cognate adaptors, while *E. coli* DnaA could not recognize either DnaD homolog (**Fig 6E**). Therefore, the interaction between DnaA and DnaD is highly specific despite the fact that it relies on a conserved interaction hot spot in domain I.

## Discussion

We have identified the protein interaction domains required to recruit and load the replicative helicase during initiation. The results indicate that most of the initiation proteins interact with the chromosome using their C-terminal ends and with their protein partners using their N-terminal ends (**Fig 7**). This is with the notable exception of the DnaB adaptor, which can bind both DNA and its protein partners with its C-terminal domain (**Figs 2C and 2D**). It was previously noted that truncating the C-terminal end of DnaB prevents it from being recruited to the origin and is lethal *in vivo* (33), which underscores the functional importance of this region.

**Figure 7.**
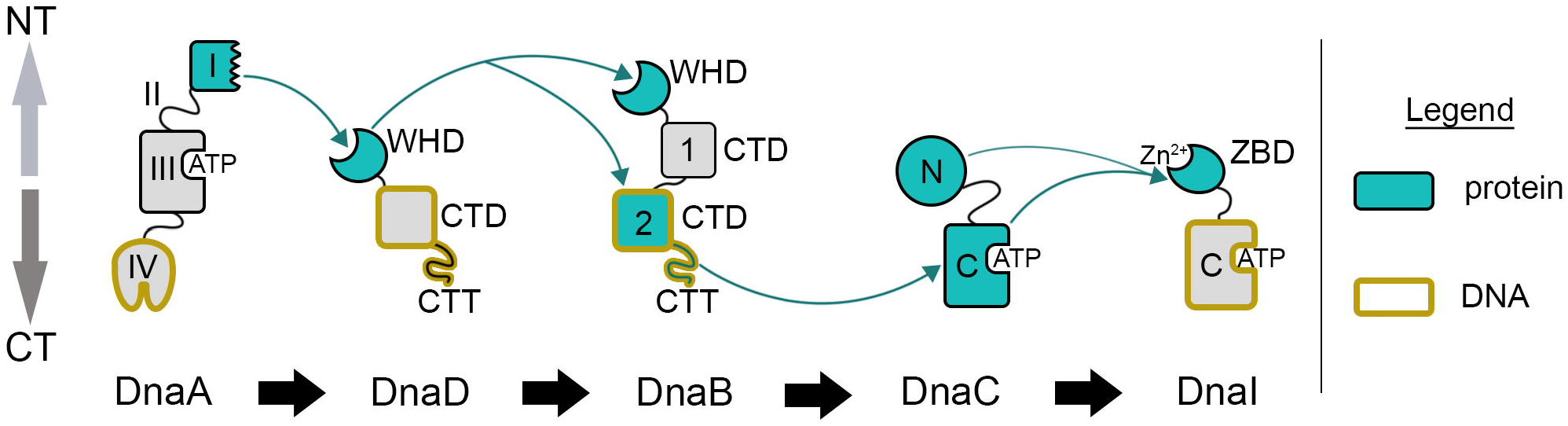
Model of Protein and DNA Interacting Domains Used for Initiation. Protein binding domains are colored in blue with interactions marked by curved blue arrows, while DNA binding domains are outlined in yellow. Domains are labeled according to Figure 2A and oriented with the N-terminal ends pointed up (NT) and the C-terminal ends pointed down (CT).

We have also mapped the interaction surfaces in the DnaA-DnaD complex using a B2H assay. This interaction required a hydrophobic cleft in the DnaD winged helix domain (**Fig 3A**), which is usually used to bind DNA in winged helix folds (34).

Interestingly, a DnaD-like plasmid initiator from *S. aureus* does use this cleft to bind DNA; however, in this case the cleft is positively charged and adopts a different conformation than the same site in *B. subtilis* DnaD (14). Therefore, the winged helix cleft is critical for DnaD-like proteins, but can be adapted for different functions. These results strongly echo licensing factors that load the replicative helicase in Eukaryotes and Archaea, which often contain winged helix domains that can bind to either DNA, protein, or both (35).

DnaD binds to an interaction hot spot on DnaA domain I and, critically, mutations on this surface that are not tolerated *in vivo* also knock out the interaction with DnaD (**Fig 5A**). Therefore, we propose that this surface is essential *in vivo* because it recruits DnaD to the origin. It is also likely that SirA competes with DnaD as part of its inhibitory function. This would prevent DnaD recruitment by DnaA and thereby ensure that the origin cannot fire during sporulation. SirA also destabilizes the interaction between DnaA and *oriC in vivo* (5), which could be a consequence of displacing DnaD.

It is intriguing that the full-length DnaA, DnaD, and DnaB proteins did not interact in the B2H, yet their isolated domains could readily associate (**Figs 2C and 2D**). We propose that these complexes are regulated through conformational changes that expose cryptic protein interaction sites, which are occluded in the full-length proteins but revealed in the isolated domains. For both DnaA and DnaD, the interacting surfaces necessary for the B2H signal were also required *in vivo* [**Fig 4A** and (8)], demonstrating these surfaces are indeed functional in the context of the full-length proteins. Intriguingly, for the DnaI loader it was previously shown that the full-length protein readily interacts with its protein partner (DnaC helicase), but its DNA binding activity is cryptic (22). Regardless of which activity is masked, our results suggest that factors which lead to conformational changes may play a critical role in regulating initiation frequency in *B. subtilis*.

## Materials and Methods

### Plasmid Construction

All plasmids were constructed using the Gibson assembly method (36) with the primers described in **S1 Table**. The assembled plasmids were isolated from single colonies and sent to the University of Michigan Sequencing Core for verification (https://seqcore.brcf.med.umich.edu/).

Plasmids used for the B2H assays are described in **S2 Table** including a list of all the primers used for Gibson assembly. The vectors were amplified using the following primers: pUT18C (oLM140 and oLM141), pUT18 (oLM218 and oLM217), pKT25 (oLM146 and oLM147), and pKNT25 (oLM212 and oLM211).

Plasmids used for integrating *dnaD* variants at the *amyE* locus are described in **S3 Table** including a list of all the primers used for Gibson assembly. All *dnaD* gene variants were cloned into pJS103 (37) which has a xylose inducible promoter driving expression of *dnaD* linked to an erythromycin resistance cassette. These regions are flanked by sequences homologous to *amyE* which allows for the plasmid to integrate at this locus through a double-crossover recombination event. The pJS103 vector was amplified by oLM197 and oLM198 in preparation for Gibson assembly.

Plasmids used for protein expression are described in **S4 Table** including a list of all the primers used for Gibson assembly. The gene fragment encoding the DnaD^WHD^ (amino acids 1-128) and its variants as well as DnaB^WHD^ (amino acids 1-153) were cloned into pET28a encoding an N-terminal HIS_6_-MBP-SUMO tag (38). The vector was amplified with oJS638 and oJS639 in preparation for Gibson assembly. The gene fragments encoding DnaD^CTD^ (amino acids 129-232), DnaB^CTD1^ (amino acids 185-300) and DnaB^CTD2^ (amino acids 303-472) were cloned into pE-SUMO. The pE-SUMO vector was amplified using oLM1 and oLM2. Note that digesting the tag with Ulp1 protease will leave the native N-terminus of the proteins.

All site-directed mutagenesis was conducted using the primer extension method (39). This required the use of at least four primers: two flanking primers that amplify the desired region and two partially complementary primers that encode the desired mutation. Each flanking primer was paired with the appropriate mutagenic primer to amplify the gene in two pieces. These amplification products were then gel purified, mixed in equal molar amounts, and used as the template in a second PCR that was set with the flanking primers only.

### Building the *dnaD23* strain

The *dnaD23* strain was created using CRISPR/Cas9 gene-editing to introduce an A166T mutation in *dnaD* using a method we developed previously (38,40). A detailed description of the process can be viewed in the supporting information section.

### Purification of DnaD^WHD^ and DnaB^WHD^

BL21 (DE3) cells were transformed with the relevant expression plasmid (**S4 Table**) and grown at 37°C until an OD_600_ of 0.7. Protein expression was induced using 0.5 mM Isopropyl β-D-1-thiogalactopyranoside (IPTG) and the cells were incubated at 37°C for 3 hours. The cells were then harvested using centrifugation, washed once with 1X PBS to remove residual media, and resuspended in 50 mL buffer A (20 mM TRIS pH 8.0, 300 mM NaCl, 5% glycerol, 1 mM DTT) with a protease inhibitor tablet (Complete EDTA-free, Roche). Sonication was used to lyse the sample, after which the soluble fraction was obtained through centrifugation and loaded onto a 4 mL amylose gravity column. The column was washed four times with 10 mL of buffer A, followed by eluting any bound protein using buffer A supplemented with 10 mM D-maltose while collecting 1 mL fractions.

Fractions containing the desired protein were pooled and adjusted to 100 mM NaCl prior to digesting the HIS_6_-MBP-SUMO tag with Ulp1 protease for two hours at room temperature. The digested sample was filtered and loaded onto a 5 mL HiTrap^TM^ Q FF column (GE life sciences) equilibrated with buffer B (20 mM TRIS pH 8.0, 1 mM DTT, 1 mM EDTA, 5% glycerol) supplemented with 100 mM NaCl. The protein was then eluted with a 90 minute gradient from 100 mM to 500 mM NaCl at 1 mL/min and injected into a HiPrep^TM^ 16/60 Sephacryl^TM^ S-200 HR column (GE life sciences) using buffer B containing 150 mM NaCl at 0.75 mL/min. The final eluted sample was concentrated using an Amicon centrifugal filter (Amicon Ultra-15 with Ultracel-10 membrane). The sample was supplemented with 25% glycerol and flash frozen for storage at −80°C.

### Purification of DnaD^CTD^, DnaB^CTD1^, and DnaB^CTD2^

BL21 (DE3) cells were transformed with the relevant expression plasmid (**S4 Table**) and grown at 37°C until an OD_600_ of 0.7. Protein expression was induced using 0.5 mM IPTG and the cells were incubated at 12°C for 16 hours. The cells were then harvested using centrifugation, washed once with 1X PBS to remove residual media, and resuspended in 50 mL buffer C (20 mM TRIS pH 8.0, 400 mM NaCl, and 5% glycerol) with a protease inhibitor tablet (Complete EDTA-free, Roche). Sonication was used to lyse the sample, after which the soluble fraction was obtained through centrifugation and loaded onto a 5 mL HiTrap^TM^ IMAC FF column (GE life sciences) equilibrated with Buffer C. The column was washed with 40 mL of Buffer C supplemented with 2 M NaCl followed by 30 mL of Buffer C with 15 mM imidazole. The bound protein was then eluted using Buffer C supplemented with 300 mM imidazole while collecting 1 mL fractions. Fractions containing the desired protein were pooled and added dropwise to 50 mL of Buffer D (20 mM pH 8.0, 1 mM DTT, 5% glycerol, and 150 mM NaCl). The tag was digested using Ulp1 protease for 2 hours at room temperature.

The digested sample was dialyzed overnight against Buffer C at 4°C. The sample was then filtered and loaded onto a 5 mL HiTrap^TM^ IMAC FF column equilibrated with Buffer C. The column was washed with 15 mL of Buffer C followed by 15 mL of Buffer C supplemented with 15 mM imidazole. The washes that contained the tagless protein were pooled and concentrated using an Amicon centrifugal filter (Amicon Ultra-15 with Ultracel-10 membrane). The concentrated protein was injected into a HiPrep^TM^ 16/60 Sephacryl^TM^ S-200 HR column (GE life sciences) using buffer E (20 mM TRIS pH 8.0, 400 mM NaCl, 1 mM DTT, 1 mM EDTA, and 5% glycerol) at 0.75 mL/min. The final eluted sample was concentrated using an Amicon centrifugal filter (Amicon Ultra-15 with Ultracel-10 membrane), supplemented with 25% glycerol, and flash frozen for storage at −80°C.

### Bacterial two-hybrid assays

BTH101 cells were co-transformed with a plasmid encoding a T18 fusion of interest and a plasmid encoding a T25 fusion of interest (details of the specific plasmids used can be viewed in **S2 Table**). Co-transformants were grown in 3 mL of LB media (supplemented with 100 μg/mL ampicillin and 25 μg/mL kanamycin) at 37°C until an OD_600_ of between 0.5 and 1.0. The cultures were adjusted to an OD_600_ of 0.5, diluted 1/1000 in LB, and spotted (5 μL per spot) onto LB agar plates containing 40 μg/mL of X-Gal (5-bromo-4-chloro-3-indoxyl-β-D-galactopyranoside), 0.5 mM IPTG, 100 μg/mL ampicillin, and 25 μg/mL kanamycin. The plates were incubated for two days at 30°C followed by an additional 24 hours at room temperature while being protected from light. All two-hybrid experiments were performed a minimum of three times working from fresh co-transformations.

### Chemical Cross-linking of DnaD^WHD^

Purified DnaD^WHD^ and its variants (16 μM final concentration) were mixed with increasing amounts of glutaraldehyde (0.03%, 0.06%, 0.12%) in 10 μL reactions containing 20 mM HEPES pH 7.7, 150 mM NaCl, 1 mM DTT, 1 mM EDTA, and 5% glycerol. Each reaction was incubated for 15 minutes at room temperature, followed by quenching with 1 μL of 1 M TRIS pH 8.0 and incubating an additional 10 minutes. SDS-loading dye (62.5 mM TRIS pH 6.8, 10% glycerol, 2% SDS, 100 mM DTT, and 0.01% bromophenol blue) was then added to the samples, which were loaded onto a 4-20% gradient SDS-polyacrylamide gel (Bio-Rad), electrophoresed in 1X TAE running buffer for 30 minutes at 200 V, and stained with coomassie blue.

### Complementation of *dnaD23ts* by DnaD Mutants

The relevant strains (**S5 Table**) were streaked onto LB agar containing 0.001% D-xylose and incubated at 30°C (permissive temperature) or 48°C (restrictive temperature) overnight. To test for the small colony phenotype, strains were instead plated on LB agar with (1%) or without (0%) D-xylose and incubated overnight at 30°C.

### Modeling the DnaA-DnaD Complex

The structures of DnaA^DI^ (PDB 4TPS) and DnaD^WHD^ (PDB 2V79) were used to model the interaction between the DnaA^DI^ ***α***2− ***α***3 surface and the DnaD^WHD^ winged helix cleft (8,20). The final model was generated using the Rosetta Docking program available on ROSIE (Rosetta Online Server that Includes Everyone) (29–31). Figures were prepared using the PyMOL Molecular Graphics System (v1.8.0.3), Schrödinger, LLC.

## Acknowledgments

This work was supported by R01GM107312 to L.A.S. We wish to thank Jeremy Schroeder for contributing the pJS103 plasmid and Peter Burby for contributing the pPB41 plasmid used for CRISPR/Cas9 gene editing in *B. subtilis*.

## Supporting Information

S1 Fig. Detecting Interactions Between DnaA, DnaD and DnaB Using Minimal Media with Maltose.

B2H of T18-tagged DnaD domains co-expressed with either T25-tagged DnaA domains (A) or T25-tagged DnaB domains (B) grown on LB or minimal media supplemented with 0.2% D-maltose (MM + Maltose). Red boxes indicate positive interactions.

S2 Fig. The DnaD^WHD^ NT Extension is Not Necessary for Protein Interactions.

(A) Schematic of DnaD^WHD^ showing the N-terminal extension (residues 1-18) that mediates tetramerization. (B) Size exclusion chromatogram of either wild type DnaD^WHD^ (black) or the ΔNT variant (blue) with elution volume (mL) labeled on the x-axis and absorbance at 280 nm (A_280_) labeled on the y-axis. The ΔNT variant elutes later than expected for the tetramer (size of the ΔNT tetramer is 51.9 kDa) demonstrating that this truncation has shifted the equilibrium to favor the dimer. (C) B2H of T18-tagged wild type DnaD^WHD^ or the ΔNT variant co-expressed with T25-tagged DnaA^DI^, DnaB^WHD^ or DnaB^CTD2^.

S3 Fig. Detecting Interactions with the DnaC Helicase Using Minimal Media with Maltose.

B2H of T18-tagged DnaC domains co-expressed with either T25-tagged DnaI domains (A) or T25-tagged DnaB domains (B) grown on LB or minimal media supplemented with 0.2% D-maltose (MM + Maltose). Red boxes indicate positive interactions.

S1 Table. Primers Used to Construct Plasmids.

S2 Table. Plasmid List for B2H Assay.

S3 Table. Plasmids Used to Integrate *dnaD* Variants at the *amyE* Locus.

S4 Table. Plasmids Used for Protein Purifications.

S5 Table. *B. subtilis* Strains Used in this Study.

